# The beneficial effect of chronic muscular exercise on muscle fragility is increased by *Prox1* gene transfer in dystrophic *mdx* muscle

**DOI:** 10.1101/2021.07.22.453435

**Authors:** Alexandra Monceau, Clément Delacroix, Mégane Lemaitre, Gaelle Revet, Denis Furling, Onnik Agbulut, Arnaud Klein, Arnaud Ferry

**Author notes:** Correspondance: A. Ferry, Centre de Recherche en Myologie, UMRS974, G.H. Pitié-Salpétrière, 47, bld de l’Hôpital, 75651 Paris cedex 13, France.

## Abstract

**Purpose:** Greater muscle fragility is thought to cause the exhaustion of the muscle stem cells during successive degeneration/repair cycles, leading to muscle wasting and weakness in Duchenne muscular dystrophy. Chronic voluntary exercise can partially reduce the susceptibility to contraction induced-muscle injury, i.e., muscle fragility, as shown by a reduced immediate maximal force drop following lengthening contractions, in the dystrophic *mdx* mice. Here, we studied the effect of Prospero-related homeobox factor 1 gene (*Prox1*) transfer (overexpression) on fragility in chronically exercised *mdx* mice, because *Prox1* promotes slower type fibres in healthy mice and slower fibres are less fragile in *mdx* muscle.

**Methods:** *mdx* mice received or not *Prox1* transfer into the *tibialis anterior* muscle and performed voluntary running into a wheel during 1 month. We also performed *Prox1* transfer in sedentary *mdx* mice. *In situ* maximal force production of the muscle in response to nerve stimulation was assessed before, during and after 10 lengthening contractions. Molecular and cellular parameters were also evaluated.

**Results:** Interestingly, *Prox1* transfer reduced the force drop following lengthening contractions in exercised *mdx* mice (p < 0.05 to 0.01), but not in sedentary *mdx* mice. It also increased the muscle expression of *Myh7* (p < 0.001), MHC-2x (p < 0.01) and *Trpc1* (p < 0.01), whereas it reduced that one of *Myh4* (p < 0.001) and MHC-2b (p < 0.01) in exercised *mdx* mice. Moreover, *Prox1* transfer decreased the maximal force (p < 0.01) before lengthening contraction in exercised *mdx* mice (p < 0.01), and reduced muscle weight (p < 0.0001) despite increased *Mstn* expression (p < 0.001).

**Conclusion:** Our results indicate that the beneficial effect of *Prox1* transfer on muscle fragility is only observed in chronically exercised *mdx* mice. Thus, *Prox1* transfer combined to chronic exercise have the potential to substantially slow the progression of the dystrophic disease in the long term.

## Introduction

Duchenne muscular dystrophy (DMD), the most common X-linked inherited muscular disease, is caused by mutations in the *DMD* gene, leading to dystrophin deficiency that results in skeletal muscle fibre injury and progressive muscle wasting and weakness. Dystrophin is a costameric protein that plays a role in force transmission and sarcolemma stability in skeletal muscle (1). In line, muscle of dystrophin-deficient *mdx* mouse, the “classic” animal model for DMD, exhibits two important functional dystrophic features. First, muscular weakness that is the decrease of the specific maximal force (the absolute maximal force generated relatively to muscle cross-sectional area or weight) with an unmodified/maintained absolute maximal force due to the muscle hypertrophy (2). Second, muscle fragility that is revealed by the high susceptibility of the fast and low oxidative *mdx* muscle for damages caused by the lengthening (eccentric) contractions, leading to an immediate marked force drop following lengthening contractions (2). This force drop is proportional to both the length of the stretch and the absolute maximal lengthening force produced during the first contraction in fast *mdx* muscle (3,4). The greater fragility in *mdx* mice is associated to reduced muscle excitability (5–8), and several genes coding ion membrane channels interacting with dystrophin are involved in muscle excitability, such as *Scn4a, Cacna1s, Slc8a1, Trpc1* and *chrna1* (9,10). Increased fragility is also related to NADPH oxidase 2 (NOX2) activity (11–13) and aggravated by inactivation of *Utrn* and *Des* coding utrophin and desmin respectively in *mdx* mice (14,15).

The fragility of the dystrophic muscle is thought to cause the exhaustion of the muscle stem cells during successive degeneration/repair cycles (16). Thus, attempt to reduce this fragility is very important because it has the potential to slow the progression of the dystrophic disease. Interestingly, chronic muscular exercise can improve (reduced) the fragility in *mdx* mice (8,17–19). In particular, voluntary running decreases fragility, i.e., reduces the force drop following lengthening contractions, in *mdx* mouse fast muscle (8,18), whereas physical inactivity aggravates it (18). Recently, it was found that the reduced fragility induced by voluntary running in *mdx* mice was related to calcineurin pathway activation, and changes in the program of genes involved in slower contractile features of muscle fibre and genes coding membrane ions channels involved in muscle excitability (8). However, voluntary running only partly reduced the susceptibility to exercise-induced muscle injury (8,18), so it would be interesting to combined the effects of exercise with those of another treatment.

While voluntary exercise offers potential therapeutic benefit, additional adjunct therapies could further improve functional dystrophic features. In the recent years, genetic or pharmacological treatments promoting slower and more oxidative fibres are been shown to be beneficial in the *mdx* mice. In fact, several studies support the idea that activation of the AMPK, calcineurin, E2F1, ERRγ, IGF1, SIRT1 and PGC1 signalling pathways alleviates some of the dystrophic features in *mdx* muscle (20–29). For example, genetic activation of calcineurin pathway improves fragility in fast muscle of the *mdx* mouse, but decreases maximal force production, thus, aggravating weakness (27). Recently, in healthy fast muscle, it was demonstrated that the loss of Prospero-related homeobox factor 1 (*Prox1*), a transcription factor essential for the development of several organs and highly conserved among vertebrates, represses the expression of slow contractile genes, whereas its overexpression via *Prox1* transfer has the opposite effect and down-regulates the fast contractile genes, (30,31). In particular, the inactivation of *Prox1* reduces the expression of the slowest myosin heavy chain *Myh7* in fast healthy muscle, without affecting oxidative capacity (succinate dehydrogenase staining) and absolute maximal force (31). *Prox1*, that is more expressed in slow fibres, is involved in the activation of the NFAT/calcineurin pathway, and promotes the slow contractile gene program in healthy muscle (30).

The principal purpose of the present study was to determine whether *Prox1* transfer, i.e., *Prox1* overexpression, reduced muscle fragility in voluntary exercised *mdx* mice. The study includes physiological outcome measurement of fragility, complemented by cellular and molecular analyses. Because we found that voluntary running and *Prox1* transfer have additive beneficial effects on fragility, a second set of experiment was performed to compared the effect of *Prox1* transfer on fragility between voluntary and sedentary *mdx* mice. Interestingly, *Prox1* transfer only reduced fragility in voluntary exercised *mdx* mice, but not in sedentary *mdx* mice.

## Materials and Methods

### Animal groups and voluntary running

All procedures were performed in accordance with national and European legislations and were approved by our institutional Ethics Committee “Charles Darwin” (Project # 01362.02). Male mice with exon 23 mutation in the *dmd* gene encoding dystrophin (Mdx mice) were used (hybrid background C57Bl/6 x C57Bl/10). Mice (2-3 months of age) were randomly divided into different control and experimental groups (Figure 1). In the first set of experiment, Mdx mice were placed (Mdx+W) in separate cages containing a wheel and were allowed to run 1-month ad libitum. The muscles of Mdx mouse runners received (Mdx+W+P) or not (Mdx+W) *Prox1* transfer into the muscle 3 days before the initiation of voluntary exercise. The running distances were collected and daily running distance was 4.2 ± 0.1 km/day. A group of sedentary Mdx mice was also studied (Mdx). The first set of experiment was performed to study the effect of *Prox1* transfer in exercised Mdx mice. Because we found an effect of *Prox1* transfer on fragility in Mdx+W+P muscle, we then performed a second set of experiment to compare the effect of *Prox1* transfer on fragility between exercised muscle and sedentary muscle. In the second set of experiments, the muscles of sedentary Mdx mice received (Mdx+P) or not (Mdx) *Prox1* transfer. The muscles were measured and collected 4 weeks after *Prox1* transfer.

**Figure 1.**
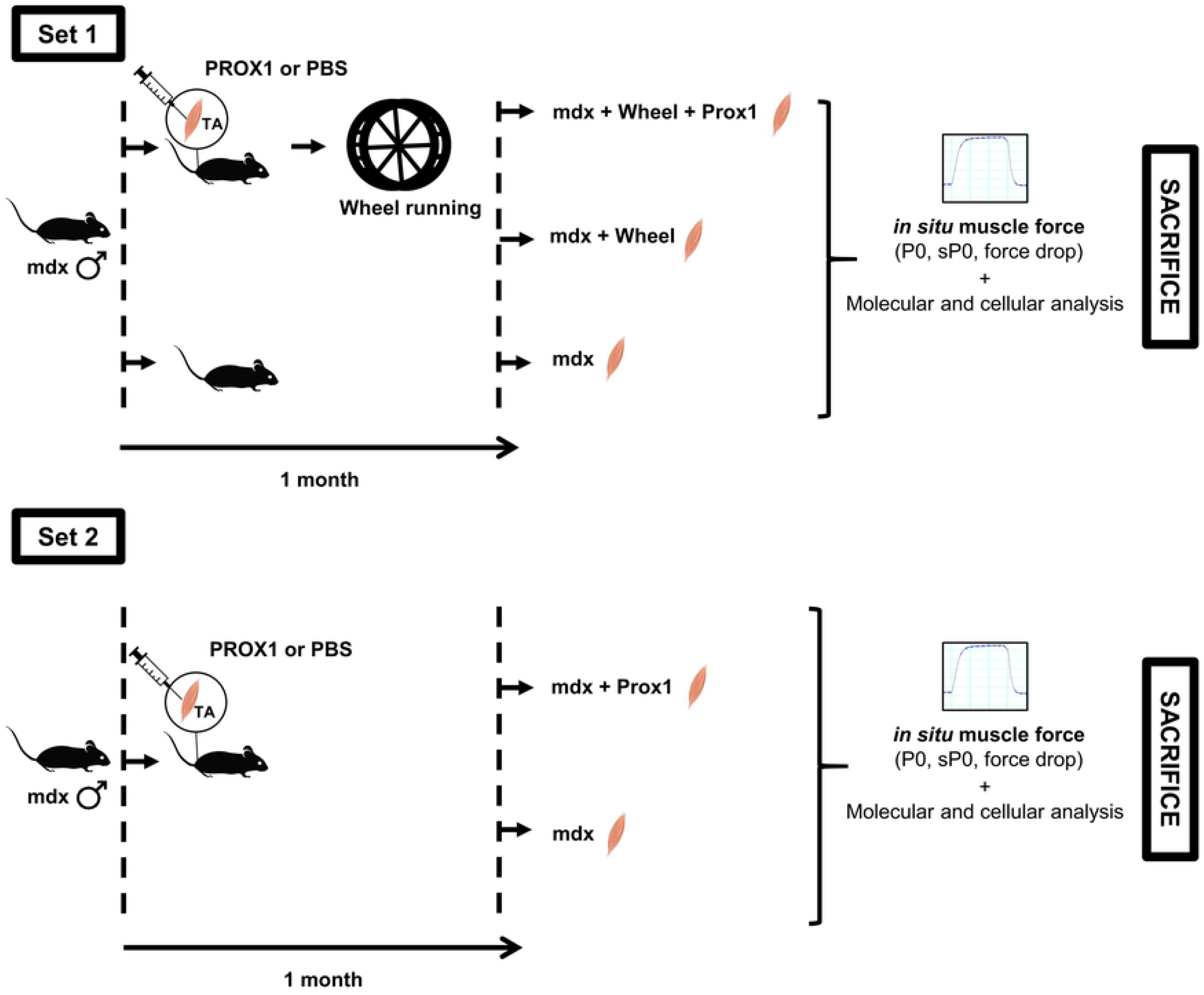
Experimental design. Two sets of experiments were performed. In the first set of experiment, we want to determine the effect of *Prox1* transfer on fragility in voluntary exercised mdx mice. The second set of experiment was to compare the effect of *Prox1* transfer between voluntary exercised and sedentary mdx mice.

### *Prox1* transfer

To overexpress Prox1, adeno-associated vectors (AAV9) carrying the *Prox1* construct (AAV-Prox1)(30) was injected in one of the Tibialis anterior (TA) muscles of the mouse (2.1 x 10^11^ vector genomes). The other TA muscle (control muscle) was injected with saline solution only. The mouse was anesthetized (3% isoflurane) and TA muscles were injected (30 μl). Briefly, hProx1 cDNA’s were cloned into psub plasmid (promoter CMV)(30). The plasmid was purified using the PureYield™ endotoxin-free Plasmid Maxiprep System (Promega, Lyon, France) and then verified by restriction enzyme digestion and by sequencing (Eurofins MWF Operon, Ebersberg, Germany). The AAV-Prox1 was produced in human embryonic kidney 293 cells by the triple-transfection method using the calcium phosphate precipitation technique. The virus was then purified by 2 cycles of cesium chloride gradient centrifugation and concentrated by dialysis. The final viral preparations were kept in PBS solution at −80°C. The number of viral genomes was determined by a quantitative PCR. Titer for AAV-Prox1 was 7.1 x 10^12^ vector genomes (vg).ml^-1^.

### Muscle fragility measurement

Muscle fragility (susceptibility to contraction-induced injury) was evaluated by measuring the *in situ* TA muscle contraction properties in response to nerve stimulation, as described previously (5). Fragility was estimated from the force drop resulting from lengthening contraction-induced injury. Briefly, mice were anesthetized using pentobarbital (60 mg/kg, ip). Body temperature was maintained at 37°C using radiant heat. The knee and foot were fixed with pins and clamps and the distal tendon of the muscle was attached to a lever arm of a servomotor system (305B, Dual-Mode Lever, Aurora Scientific) using a silk ligature. The sciatic nerve was proximally crushed and distally stimulated by a bipolar silver electrode using supramaximal square wave pulses of 0.1 ms duration. We first determined the optimal length (L0, length at which maximal isometric force was obtained during the tetanus). Once L0 was obtained, a maximal isometric contraction of the TA muscle was initiated during the first 500 ms. Then, muscle lengthening (10% L0) at a velocity of 5.5 mm/s (0.85 fibre length/s) was imposed during the last 200 ms. Nine lengthening contractions of the TA muscles were performed in Mdx mice, each separated by a 60 s rest period. Absolute maximal isometric force was measured 1 min after each lengthening contraction and expressed as a percentage of the initial maximal force (force drop). Absolute maximal isometric force measured before the first lengthening contraction was also normalized to the muscle mass in order to calculate the specific maximal isometric force, an index of muscle weakness. In addition, we measured the absolute maximal lengthening force during the first lengthening contraction, and index of the muscle stress. After contractile measurements, the animals were killed with cervical dislocation.

### Real-time quantitative PCR (polymerase chain reaction)

Muscles (TA) were snap frozen in liquid nitrogen and stored at −80°C until use. Total RNA was isolated from TA muscles using Trizol (Invitrogen). Complementary DNA (cDNA) was then synthesized from 1 μg of total RNA using the RevertAid First Strand cDNA Synthesis kit with random hexamers, according to the manufacturer’s instructions (Thermo Scientific). RT-PCR was performed on a LightCycler 480 System at the platform iGenSeq of the Institut du Cerveau et de la Moelle epinière, using LightCycler 480 SYBR Green I Master Mix (Roche, Basel, Switzerland)(5). The expression of *Hmbs* was used as reference transcript because it’s expression did not differ between groups. The 2-ΔΔCP method has been used as a relative quantification strategy for quantitative real-time polymerase chain reaction (qPCR) data analysis. All sequences of primers used are presented in Table 1.

**Table 1.**
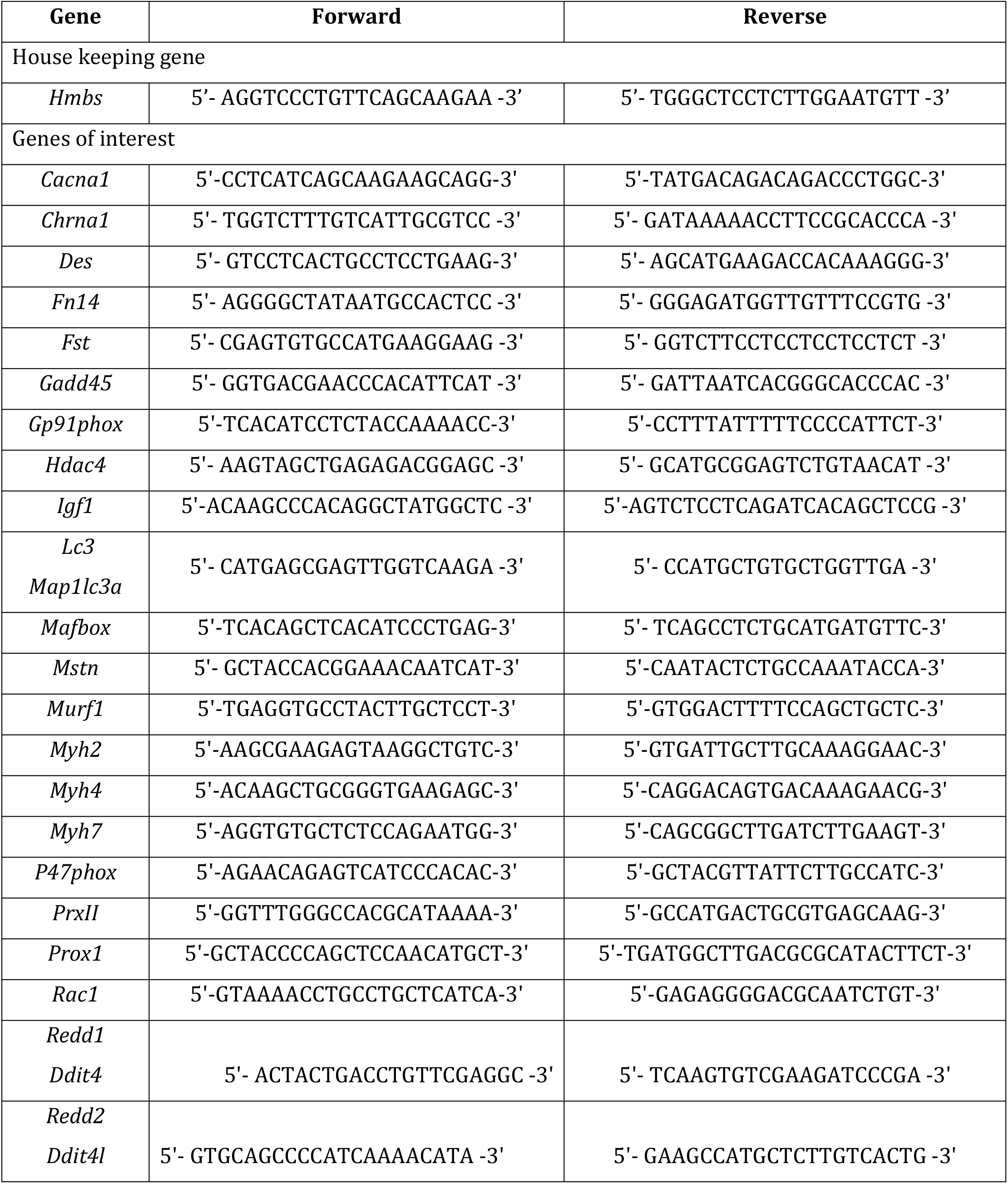
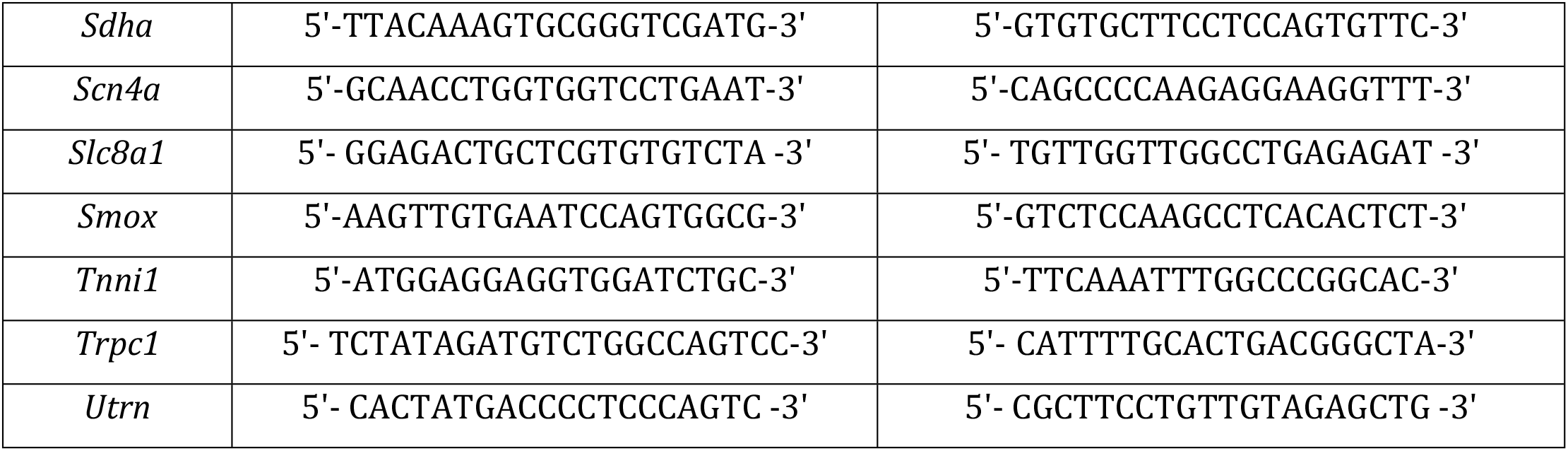
Sequences of primers used.

### Histology

Transverse serial sections (8 μm) of TA muscles were obtained using a cryostat, in the mid-belly region. Some of sections were processed for histological analysis according to standard protocols (succinate dehydrogenase, SDH, a marker of oxidative capacity). Other sections were processed for immunohistochemistry. For determination of muscle fibre diameter, fibre expressing myosin heavy chain (MHC) frozen unfixed sections were blocked 1h in phosphate buffer saline plus 2% bovine serum albumin, 2% fetal bovine serum. Sections were then incubated overnight with primary antibodies against, laminin (Sigma, France), MHC-2a (clone SC-71, Developmental Studies Hybridoma Bank, University of Iowa), MHC-2b (clone BF-F3, Developmental Studies Hybridoma Bank), MHC-1 (clone BA-D5, Developmental Studies Hybridoma Bank). After washes in PBS, sections were incubated 1 h with secondary antibodies (Alexa Fluor, Invitrogen). Slides were finally mounted in Fluoromont (Southern Biotech). Images were captured using a digital camera (Hamamatsu ORCA-AG) attached to a motorized fluorescence microscope (Zeiss AxioImager.Z1), and morphometric analyses were made using the software ImageJ. The magnification used for quantification of histological images was 20x. MHC-2x fibres were identified as fibres that do not express MHC-2b, MHC-2a (or MHC-1). The numbers of fibre type expressing MHC-1, MHC-2x and MHC-2a per muscle cross-section were determined. We attempt to analyze all the fibers of the muscle section, but some were excluded from the analysis for reasons of improper labeling. Data presented correspond to pure MHC-1, MHC-2x and MHC-2a expressing fibres (without mixed MHC coexpression).

### SDS-PAGE electrophoresis of MHC isoforms (proteins)

The muscles were extracted on ice for 60 min in four volumes of extracting buffer containing 0.3 M NaCl, 0.1 M NaH2PO4, 0.05 M Na2HPO4, 0.01 M Na4P2O7, 1 mM MgCl2, 10 mM EDTA, and 1.4 mM 2-mercaptoethanol (pH 6.5). Following centrifugation, the supernatants were diluted 1:1 (vol/vol) with glycerol and stored at - 20°C. MHC isoforms (proteins) were separated on 8% polyacrylamide gels, which were made in the Bio-Rad mini-Protean II Dual slab cell system. The gels were run for 31 h at a constant voltage of 72 V at 4°C (32). Following migration, the gels were silver stained. The gels were scanned using a video acquisition system. The relative level of MHC isoforms was determined by densitometric analysis using Image J software.

### Statistical analysis

Groups were statistically compared using the Prism software v8 (GraphPad, La Jolla, CA, USA). Data were tested for homogeneity of variance using a Brown-Forsythe test. For the first set of experiment, one-way ANOVA was used to analyze the following variables: mRNA expression, percentage of fibres expressing MHC-1, MHC2a or MHC-2b, percentage of weak SDH staining, absolute and specific maximal force, absolute maximal lengthening force, the ratio of absolute maximal lengthening force to the absolute maximal lengthening force, muscle weight, diameter of the fibres. Fragility was analyzed by two-way ANOVA, groupes (Mdx, Mdx+W, Mdx+W+P) by lengthening contractions (0, 3, 6, 9), with the repeated measures on lengthening contractions. Unpaired t-test with Welch’s correction was used to analyze the % of MHC-2x and MHC-2a (electrophoresis). For experiment 2, unpaired t-test with Welch’s correction was used for the following variable: mRNA expression, percentage of fibres expressing MHC-1, MHC2a or MHC-2b, absolute and specific maximal force, absolute maximal lengthening force, muscle weight, and diameter of the fibres. Fragility was analyzed by two-way ANOVA, groupes (Mdx, Mdx+P) by lengthening contractions (0, 3, 6, 9), with the repeated measures on lengthening contractions. Moreover, when significant main effect (ANOVA) was observed, multiple-comparisons were performed with Tukeys test. Finally, when significant interaction was found (ANOVA), differences were tested with Holm-Sidak test. Values are means ± SEM.

## Results

### *Prox1* transfer in voluntary exercised Mdx muscle promotes slower contractile features

In the first set of experiment, we first determined whether *Prox1* transfer increased slower contractile features in voluntary exercised Mdx mice. *Prox1* transfer into the TA muscle markedly increased the expression of *Prox1* (x 37.0) in voluntary exercised Mdx TA muscle (Mdx+W+P) as compared to voluntary exercised Mdx TA muscle (Mdx+W)(p < 0.0001)(Figure 2A), as assessed by qPCR analysis. We also found that the expression of *Myh7* coding for MHC-1 (x 15.1)(p < 0.001) was increased in Mdx+W+P muscle as compared to Mdx+W muscle, whereas that of *Myh4* coding for MHC-2b was reduced (x 0.6)(Figure 2B)(p < 0.001). In agreement, using gel electrophoresis technique, we found that the relative amounts (percentage of total) of MHC-2b protein were reduced (x 0.8, p < 0.01) whereas that of MHC-2x protein was increased (x 1.6, p < 0.01), respectively (Figures 1C) in Mdx+W+P muscle as compared to Mdx+W muscle. Immunohistological analyses revealed that these changes were not associated with the modification in the percentages of MHC-1, MHC-2a and MHC-2x expressing fibres because they were not different between Mdx+W+P muscle and Mdx+W muscle (Figures 2D and 2E). Moreover, there was no difference between Mdx+W+P and Mdx+W muscles in the expression of a marker of oxidative capacity, *Sdha*, a gene encoding a complex of the mitochondrial respiratory chain (Figure 2B). In line, histological analyses revealed that the percentage of the cross-sectional muscle area occupied by weak succinate dehydrogenase staining (SDH) staining was not different between Mdx+W+P muscle and Mdx+W muscle (Figure 2F).

**Figure 2.**
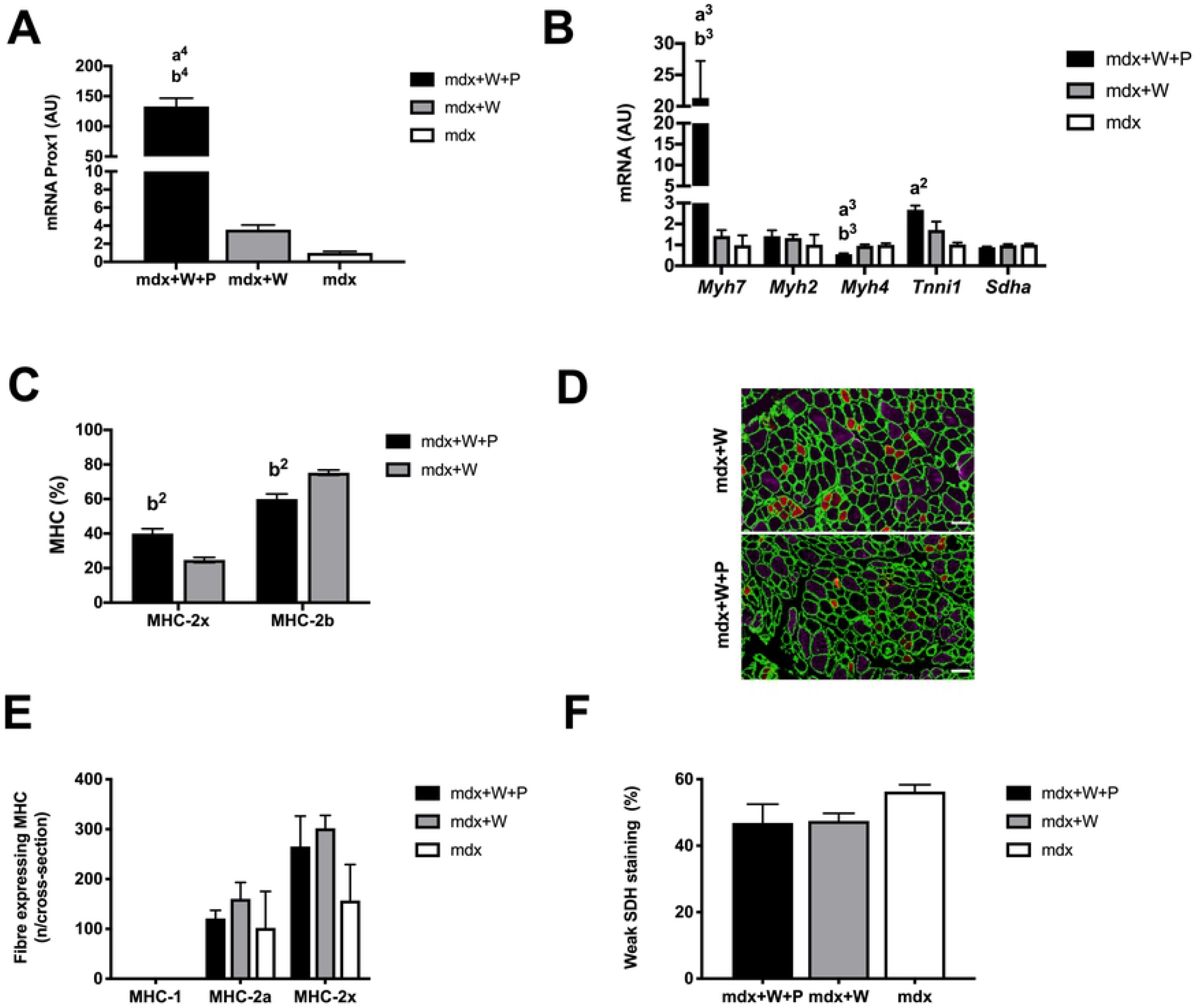
Effect of *Prox1* transfer on the expression of *Prox1* and markers of fibre type specification in voluntary exercised mdx mice (first set of experiment). (A) *Prox1* expression in Mdx+W+P and Mdx+P muscle. N = 6-8 per group. (B) Expression of genes encoding fibre type specific contractile proteins in Mdx+W+P and Mdx+P muscle. N = 6-8 per group. (C) Relative amounts of MHC-2x and MHC-2b proteins in Mdx+W+P and Mdx+P muscle. N = 3 per group. (D) Representative image of muscle cross-section of fibres expressing MHC-2a, and MHC-2b in Mdx+W+P and Mdx+P muscle. MHC-2a (red), and MHC-2b (purple) expressing fibers are labeled by specific antibodies whereas MHC-2x expressing fibres are not labeled (black). Fiber outline was visualized by antilaminin antibody (green). Less than 10 fibres expressing MHC-1 per cross-section were seen. Scale bar = 100μm. (E) Numbers per muscle cross-section of fibres expressing MHC-1, MHC-2a and MHC-2x in Mdx+W+P and Mdx+P muscle. n = 3-5 per group. (F) Percentage of the muscle cross-sectional areal occupied with weak SDH staining in Mdx+W+P and Mdx+P muscle. N = 5-8 per group. Mdx+W+P: voluntary exercised mdx muscle that received Prox1 transfer into the muscle Mdx+W: voluntary exercised mdx muscle Mdx: mdx muscle a2, a3, a4: significant different from Mdx, p < 0.01, p < 0.001, p < 0.0001, respectively. b2, b3, b4: significant different from Mdx+W, p < 0.01, p < 0.001, p < 0.0001, respectively.

These data indicate that intramuscular delivery of AAV-*Prox1* induced a substantial fast to slow contractile transition in the TA muscle of voluntary exercised Mdx mice.

### *Prox1* transfer in voluntary exercised Mdx muscle further improves muscle fragility

The first set of experiment revealed that the immediate force drop following lengthening contractions in Mdx+W muscle was reduced as compared to Mdx muscle (p < 0.0001)(Figure 3A). Interestingly, *Prox1* transfer in voluntary exercised Mdx muscle further reduced the force drop following lengthening contractions (Figure 3A). In fact, the force drops following the 6^th^ (p < 0.05) and 9^th^ (p < 0.01) lengthening contractions were lower in Mdx+W+P muscle as compared to Mdx+W muscle (Figure 3A), indicating that *Prox1* transfer improved (reduced) fragility in voluntary exercised Mdx muscle.

**Figure 3.**
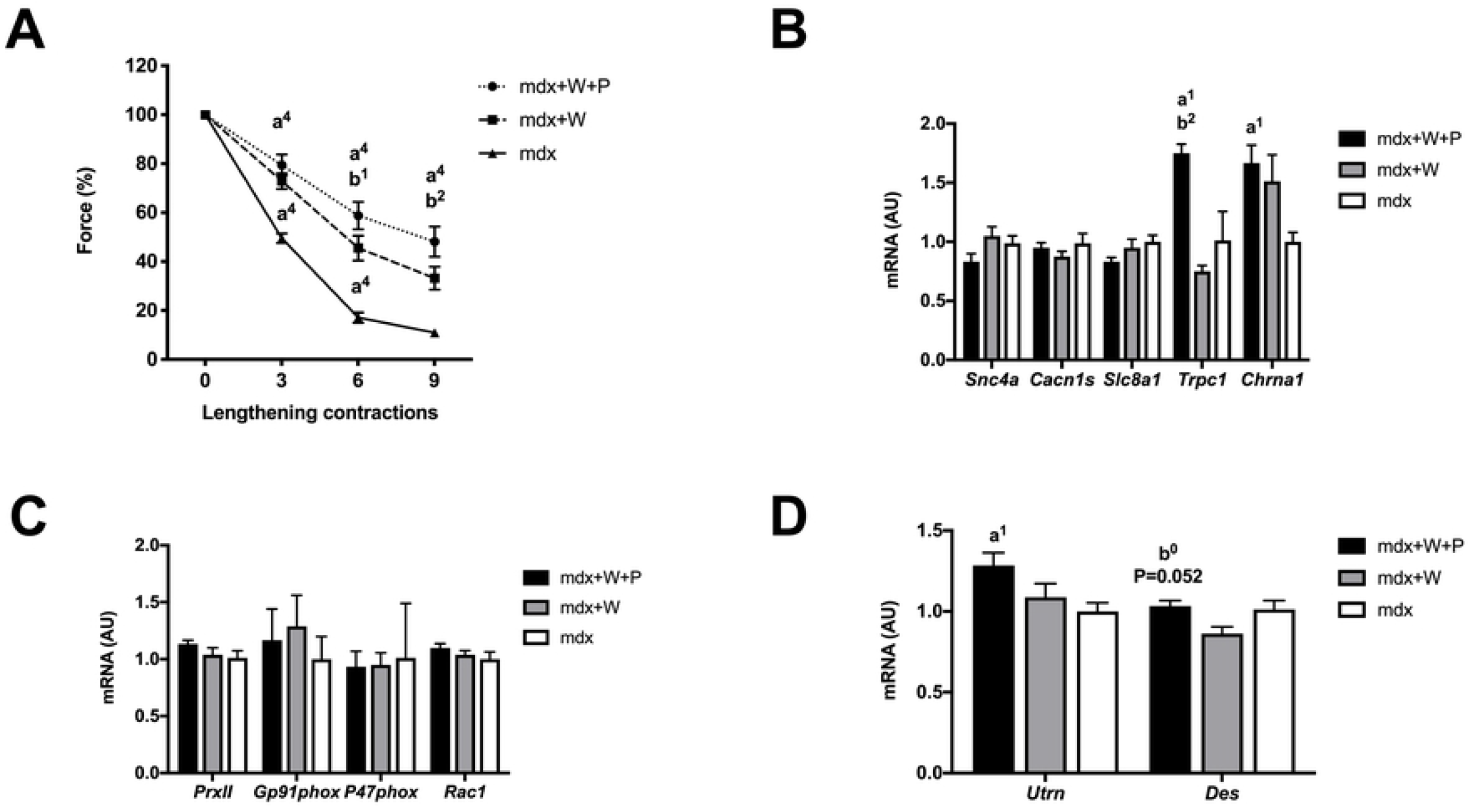
Effect of *Prox1* transfer on fragility (susceptibility to muscle injury) and related gene expression in voluntary exercised mdx mice (first set of experiment). (A) Force drop following lengthening contractions (Fragility) in Mdx+W+P and Mdx+P muscle. n=6-8 per group. (B) Expression of genes encoding ion channels, related to excitability in Mdx+W+P and Mdx+P muscle. N=6-8 per group. (C) Expression of genes, related to NADPH oxidase 2 (NOX2) in Mdx+W+P and Mdx+P muscle. N=6-8 per group. (D) Expression of genes encoding utrophin *(Utrn)* and desmin *(Des)* in Mdx+W+P and Mdx+P muscle. N=6-8 per group. Mdx+W+P: voluntary exercised mdx muscle that received Prox1 transfer into the muscle Mdx+W: voluntary exercised mdx muscle Mdx: mdx muscle a1, a4: significant different from Mdx, p < 0.05, p < 0.0001, respectively. b1, b2: significant different from Mdx+W, p < 0.05, p < 0.01, respectively.

The fast to slower contractile conversion described above can explained, at least in part, the improved fragility in Mdx+W+P muscle. Moreover, we tested the possibility that *Prox1* transfer also improved fragility via the modifications of the expression of genes coding membrane ions channels. The expression of *Trpc1* encoding for transient receptor potential cation channel subfamily C member 1 (x 2.1) was increased in Mdx+W+P muscle as compared to Mdx+W muscle (p < 0.01)(Figure 3B). No difference between Mdx+W+P and Mdx+W muscles was observed concerning the expression of *Scn4a, Cacna1s, Slc8a1 and Chrna1* (Figure 3B). Then, we determined whether the reduced force drop following lengthening contractions induced by *Prox1* transfer was associated to change (decrease) in NOX2 pathway. We found no change in the expression of *PrxII, Gp91phox, P47phox* and *Rac1* (Figure 3C) in Mdx+W+P muscle as compared to Mdx+W muscle (Figure 3C). We also determined whether *Prox1* transfer increased *Utrn* and *Des* expression. The expression of *Utrn* was not increased in Mdx+W+P muscle as compared to Mdx+W muscle, whereas that one of *Des* increased (x 1.2) in Mdx+W muscle, although not significantly (p = 0.052)(Figure 3D).

Thus, the improved TA muscle fragility induced by *Prox1* transfer in voluntary exercised mice was associated with the modification of expression of MHC-2b and MHC-2x proteins and several genes involved in different aspects of muscle function and structure (*Myh7*, *Myh4, Trpc1*).

### *Prox1* transfer in voluntary exercised Mdx muscle reduced absolute isometric maximal force and induces atrophy

The first set of experiment revealed that *Prox1* transfer combined to voluntary running and voluntary running alone did not affect specific maximal isometric force before lengthening contractions (Figure 4A). However, absolute maximal isometric force was reduced in Mdx+W+P muscle (x 0.6) as compared to Mdx+W muscle (p < 0.01)(Figure 4B), indicating that *Prox1* transfer induced muscle weakness in voluntary exercised Mdx muscle. Similarly, absolute maximal lengthening force was lower (x 0.6) in Mdx+W+P muscle (157.2 g ± 7.5) compared to Mdx+W muscle (240.0 g ± 10.8) muscle (p < 0.01). In addition, the ratio of absolute maximal lengthening force to the absolute maximal isometric force was not different between Mdx+W+P muscle (1.9 ± 0.1) and Mdx+W muscle (1.8 ± 0.1).

**Figure 4.**
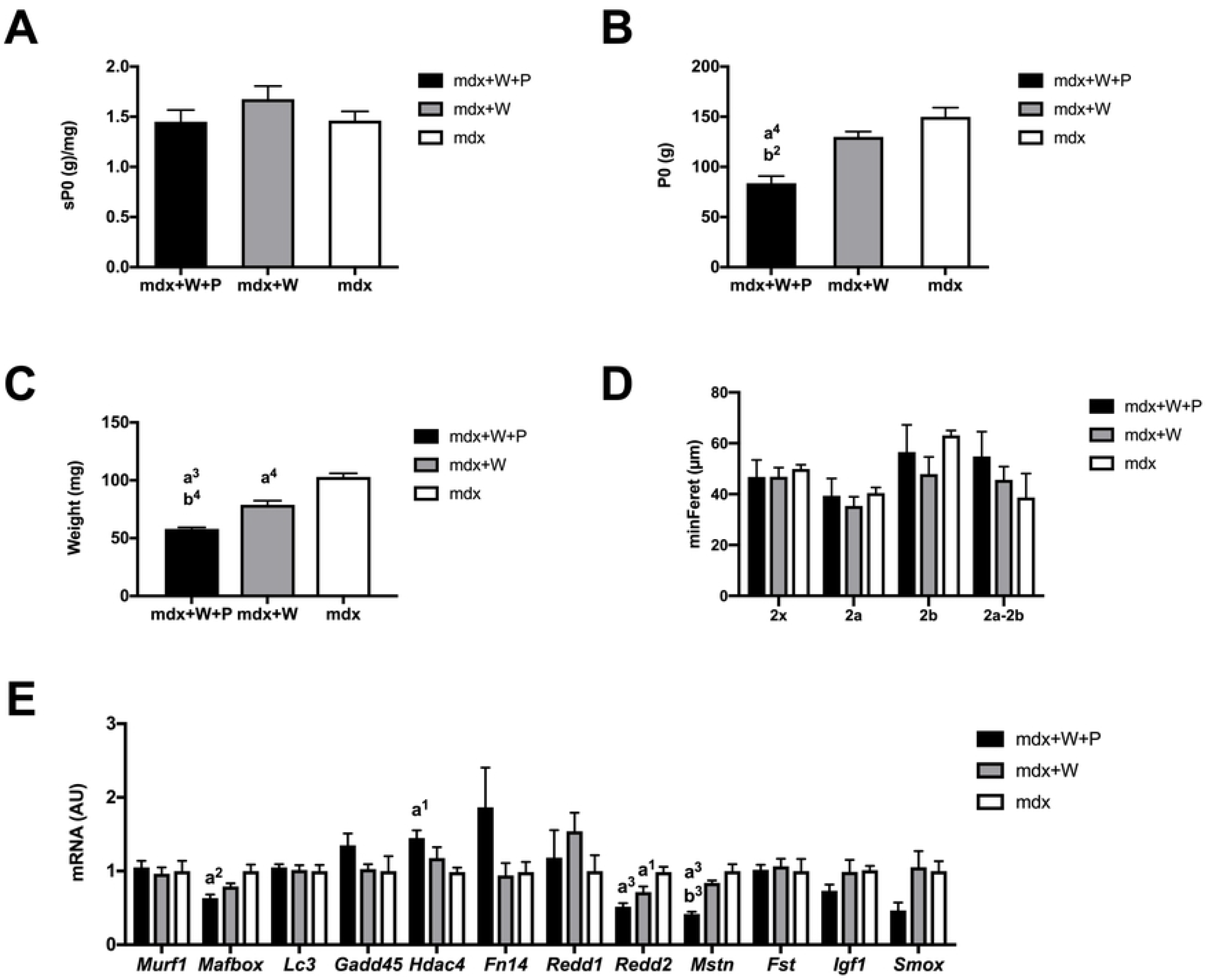
Effect of *Prox1* transfer on absolute (P0) and specific (sP0) maximal forces, muscle weight and gene expression of atrophy markers in voluntary exercised mdx mice (first set of experiment). (A) Specific maximal force in Mdx+W+P Mdx+W+P and Mdx+P muscle. n=6-8 per group (B) Absolute maximal force in Mdx+W+P and Mdx+P muscle. n=6-8 per group. (C) Muscle weight in Mdx+W+P and Mdx+P muscle. n=6-8 per group (D) Diameter (min ferret) of the fibres expressing MHC-2b, MHC-2a and MHC-2x in Mdx+W+P and Mdx+P muscle muscle. n = 4-5 per group. (E) Muscle weight in Mdx+W+P and Mdx+P muscle. n=5-9 per group (H) Diameter (min ferret) of the fibres expressing MHC-2b, MHC-2a and MHC-2x in Mdx+W+P and Mdx+P muscle. n = 4 per group. (I) Expression of genes related to atrophy in Mdx+W+P and Mdx+P muscle. N=6-8 per group. Mdx+W+P: voluntary exercised mdx muscle that received Prox1 transfer into the muscle Mdx+W: voluntary exercised mdx muscle Mdx: Mdx muscle a1, a2, a3, a4: significant different from Mdx, p < 0.05, p < 0.01, p < 0.001, p < 0.0001, respectively. b2, b3, b4: significant different from Mdx+W, p < 0.01, p < 0.001, p < 0.0001, respectively.

The reduced absolute maximal isometric force was related to a lower muscle weight (x 0.7) in Mdx+W+P muscle as compared to Mdx+W muscle (p < 0.001)(Figure 4C). The lower muscle weight was not associated with a reduced diameter of the different types of fibres (Figure 4D).

Numerous genes encoding proteins are involved in muscle growth and maintenance (33,34). The ubiquitin-proteasome system plays a key role in triggering muscle atrophy when the expressions of *Murf1* and *Mafbox* are increased. Quantitative real-time PCR revealed that the expressions of these genes were not increased in Mdx+W+P muscle as compared to Mdx+W muscle (Figure 4E). We then analyzed another atrophic mechanism, autophagy, which involves a battery of genes including *Lc3* which could contribute to the degradation of muscle proteins (35). We did not find any change in *Lc3* expression in Mdx+W+P muscle (Figure 4E). Similarly, *Gadd45, Hdac4, Fn14, Redd1, Redd2, Mstn, Fst, Igf1*, and *Smox* genes also did not seem to participate to the atrophic state of Mdx+W+P muscle (Figure 4E). For example, *Mstn*, the negative regulator of muscle growth, was down-regulated in Mdx+W+P muscle as compared to Mdx+W muscle (p < 0.001)(Figure 4E).

Thus, the reduction in maximal isometric force induced by *Prox1* transfer in voluntary exercised Mdx muscle was related to decreased muscle weight without reduction of fibre diameter, and was associated with the increased expression of *Mstn*.

### *Prox1* transfer in sedentary Mdx muscle promotes slower contractile features but does not reduce fragility

A second set of experiment was performed to compare the effect of *Prox1* transfer between voluntary Mdx mice and sedentary Mdx mice. Similarly to voluntary exercised Mdx muscle, *Prox1* transfer in sedentary Mdx muscle (Mdx+P muscle) increased the expressions of *Prox1* (x 27.3)(p < 0.0001)(Figure 5A), *Myh7* (x 6.2)(p < 0.05)(Figure 5B), reduced that one of *Myh4* (x 0.7)(p < 0.05)(Figure 5B), did not alter the percentages of MHC-1, MHC-2a and MHC-2x expressing fibres (Figures 5C), as compared to sedentary Mdx muscle. In contrast to voluntary exercised Mdx muscle, *Prox1* transfer increased the expression of *Tnni1* (x 2.4)(p < 0.01), reduced the expression of *Sdha* (x 0.8)(Figure 5B)(p < 0.01) and did not change the relative amounts of MHC-2b and MHC-2x proteins (Figures 5D) in Mdx+P muscle as compared to Mdx muscle. Overall, intramuscular delivery of AAV-*Prox1* also induced a fast to slow contractile conversion in the TA muscle of sedentary Mdx mice, but to lower extent.

**Figure 5.**
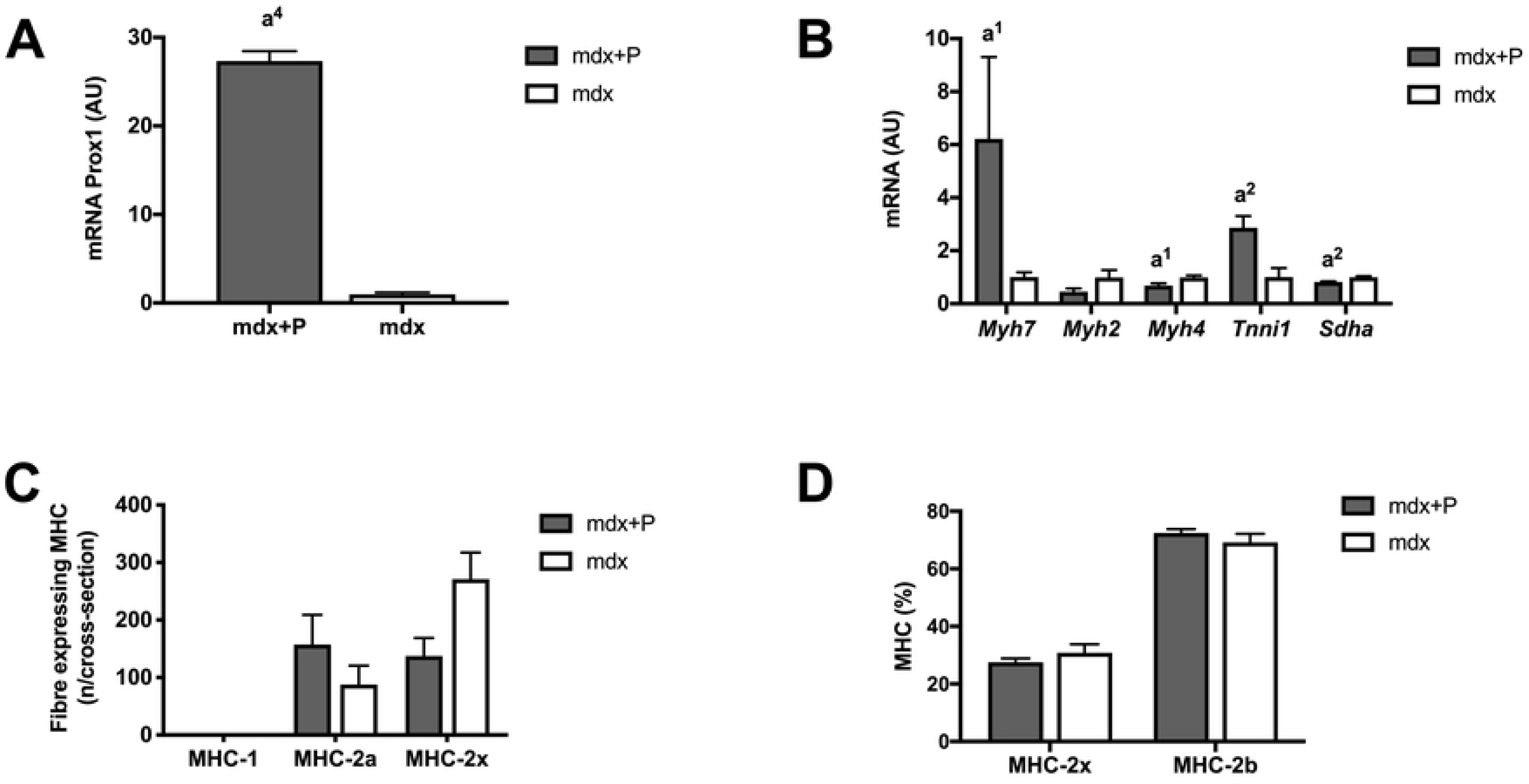
Effect of *Prox1* transfer on the expression of *Prox1* and markers of fibre type specification in sedentary mdx mice (second set of experiment). (A) Prox1 expression in Mdx+P and Mdx muscle. N = 6-11 per group. (B) Expression of genes encoding fibre type specific contractile proteins in Mdx+P and mdx muscles. N = 6-11 per group. (C) Numbers per muscle cross-section of fibres expressing MHC-1, MHC-2a and MHC-2x in Mdx+P and Mdx muscle. N= 3-5 per group. (D) Relative amounts of MHC-2x and MHC-2b proteins in Mdx+P and Mdx muscle. N = 3 per group. Mdx+P: Mdx muscle that received *Prox1* transfer into the muscle Mdx: Mdx muscle a1, a2, a4: significant different from Mdx, p < 0.05, p < 0.01, p < 0.0001, respectively.

In contrast to voluntary exercised Mdx muscle, we found in the second set of experiment that the force drop following lengthening contractions was not significantly reduced by *Prox1* transfer in sedentary Mdx muscle because there was no significant difference between Mdx+P muscle and Mdx muscle (Figure 6A). Similarly to voluntary exercised Mdx muscle, *Prox1* transfer in Mdx+P muscle increased *Trpc1* expression (p < 0.01)(Figure 6B), but to lesser extent (x 1.4), did not alter the expression of *PrxII, Gp91phox, P47phox* and *Rac1* (Figure 6C), and increased not significantly *Des* expression (Figure 6D). In contrast to voluntary exercised Mdx muscle, the expression of *Cacn1s* and *Chrna1* was increased in Mdx+P muscle compared to Mdx muscle (p < 0.01)(Figure 6B).

**Figure 6.**
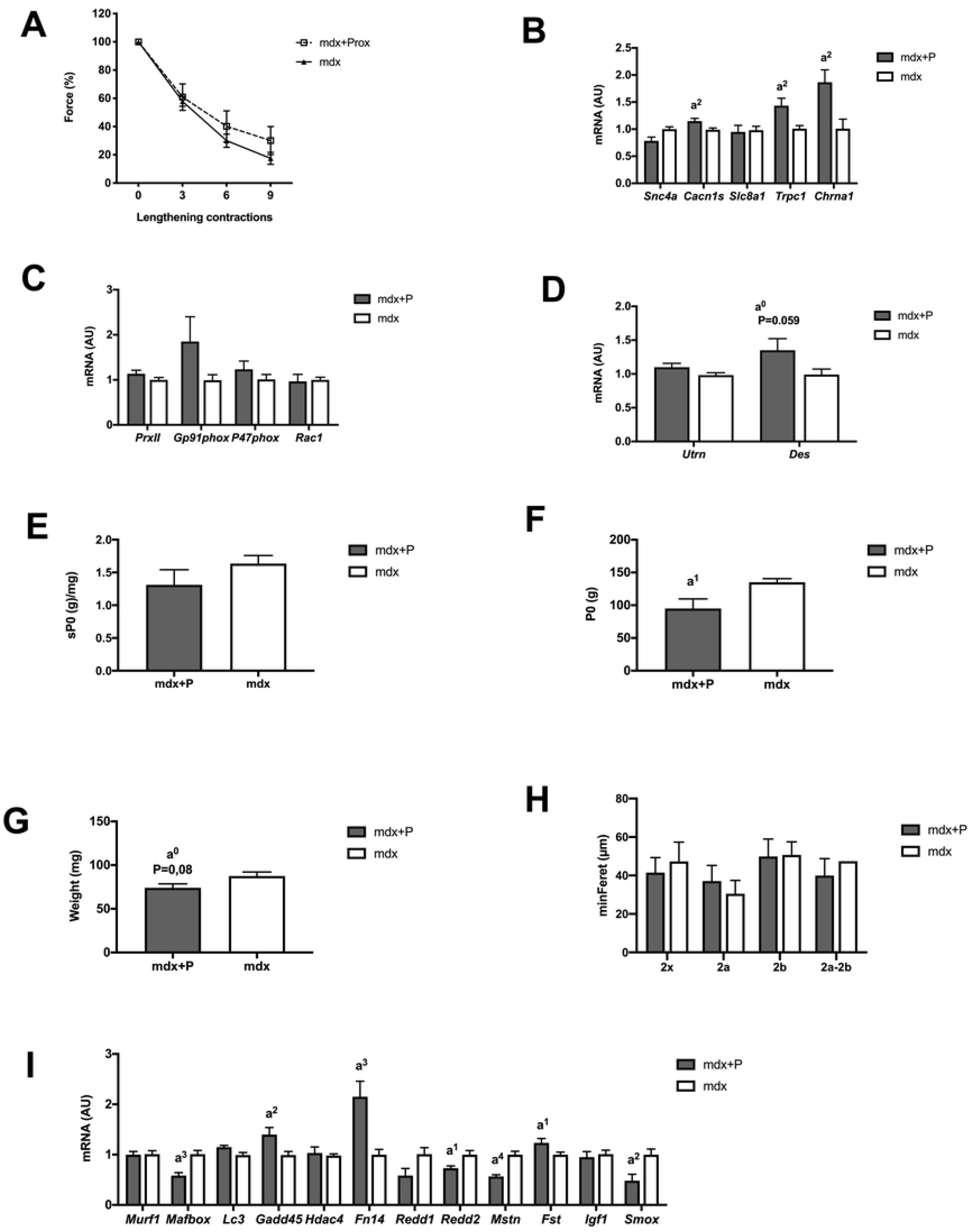
Effect of *Prox1* transfer on fragility (susceptibility to muscle injury) and related gene expression, absolute (P0) and specific (sP0) maximal forces, muscle weight and gene expression of atrophy markers in voluntary exercised mdx mice (first set of experiment) in sedentary mdx mice (second set of experiment). (A) Force drop following lengthening contractions in Mdx+P and Mdx muscle. n=5-8 per group. (B) Expression of genes encoding ion channels, related to excitability in Mdx+P and Mdx muscle. N=6-11 per group. (C) Expression of genes, related to NADPH oxidase 2 (NOX2) in Mdx+P and Mdx muscle. N=6-11 per group. (D) Expression of genes encoding utrophin (*Utrn*) and desmin (*Des*) in Mdx+P and Mdx muscle. N=6-11 per group. (E) Specific maximal force in Mdx+P and Mdx muscle. n=5-8 per group (F) Absolute maximal force in Mdx+P and Mdx muscle. n=5-8 per group (G) Muscle weight in Mdx+P and Mdx muscle. n=5-9 per group (H) Diameter (min ferret) of the fibres expressing MHC-2b, MHC-2a and MHC-2x Mdx+P and Mdx muscle. N = 4 per group. (I) Expression of genes related to atrophy in Mdx+P and Mdx muscle. N=6-11 per group. a1, a2, a3, a4: significant different from Mdx, p < 0.05, p < 0.01, p < 0.001, p < 0.0001, respectively.

Thus, *Prox1* transfer in sedentary Mdx muscle does not reduced fragility, did not change the expression of MHC-2b and MHC-2x, whereas it altered the expression of several genes involved in different aspects of muscle function (*Myh7*, *Myh4*, *Tnni1, Sdha, Trpc1, Cacn1s* and *Chrna1*).

### *Prox1* transfer in sedentary Mdx muscle also reduced absolute isometric maximal force

Similarly to voluntary exercised Mdx muscle, we found in the second set of experiment that *Prox1* transfer in Mdx+P muscle did not change specific maximal isometric force (Figure 6E), reduced absolute maximal isometric force (x 0.7)(p < 0.05)(Figure 6F), reduced muscle weight (x 0.8) although not significantly (p = 0.08)(Figure 6G), did not alter fibre diameter (Figure 6H) and decreased the expression of *Mstn* (p < 0.0001)(Figure 6I). In contrast to voluntary exercised Mdx muscle, *Prox1* transfer did not significantly (p = 0.07) decrease absolute maximal lengthening force in Mdx+P muscle (205.9 g ± 18.8) as compared to Mdx muscle (257.0 g ± 15.3). Moreover, it decreased the expression of *Mafbox* (p < 0.001), *Reed2* (p < 0.05) and *Smox* (p < 0.01), whereas it increased the expression *Gadd45* (p < 0.01), *Fn14* (p < 0.001), and *Fst* (p < 0.05) in Mdx+P muscle (Figure 6I).

Thus, *Prox1* transfer in sedentary Mdx muscle also reduced maximal isometric force production, reduced muscle weight without reduction of fibre diameter, and these effects were associated with the modifications of the expressions of *Mstn, Mafbox, Reed2, Smox, Gadd45, FN14 and Fst*.

## Discussion

### *Prox1* transfer only improves fragility in voluntary exercised Mdx mice

The present study confirms previous studies (8,18) showing that voluntary exercise alleviates the great susceptibility to contraction induced injury, a major dystrophic feature, in fast anterior crural muscles (TA and extensor digitorum longus) of *mdx* mice, such as *Dmd* based preclinical therapy (5). For the first time, we demonstrate that *Prox1* transfer further improves fragility in voluntary exercised *mdx* mice. Importantly, the muscle was protected from damaging muscle contractions by *Prox1* transfer only when *mdx* mice performed voluntary exercise. This improved fragility observed in exercised *mdx* mice treated with *Prox1* transfer might be very interesting if it is assumed that fragility causes the exhaustion of the muscle stem cells during successive degeneration/repair cycles (16). *Prox1* transfer might reduce the progressive muscle wasting in exercised dystrophic muscle because of the promotion of less fragile fibres.

This beneficial effect of *Prox1* transfer in exercised *mdx* mice could be explained by a lower work and stress during lengthening (3) because absolute maximal lengthening force (presumably work) is reduced in exercised *mdx* mice, in proportion to the absolute maximal isometric force. However, we previously found no strong association between fragility and lengthening force in *mdx* mice, when muscle is maximally activated and for a constant stretch (18). Indeed, fragility was increased by inactivity and reduced by voluntary exercise in *mdx* mice whereas absolute maximal lengthening force was respectively reduced and unchanged (18).

The reduced fragility induced by *Prox1* transfer in exercised *mdx* mice is associated with the promotion of slower contractile features (increased and reduced expression of *Myh7* and *Myh4* respectively, reduced and increased relative amounts of MHC-2b and MHC-2x proteins respectively). This relation between improved fragility and slower contractile features is in line with the 2 following points. First, slow muscle is less fragile than fast muscle in *mdx* mice (4,36). Second, exercise and pharmacological or genetic activation of signaling pathways, such as calcineurin, PPAR-β, PGC1-α, and AMPK, that promote a slower and more oxidative gene program, improve fragility in *mdx* mice (8,17,18,22,26–28,37). It was previously demonstrated that *Prox1* promotes slower features, and activates the NFAT-calcineurin pathway (30), a signaling pathway known to play an important role in fibre type specification (38).

It is also possible that *Prox1* transfer improves fragility in voluntary exercised *mdx* mice by a preserved excitability, as voluntary exercise and *Dmd* based therapy (5,8). In our experiments, reduced excitability, i.e. plasmalemma electrical dysfunction leading to defective generation and propagation of muscle potential action, largely contributes to the immediate force drop following lengthening contractions in mdx mice (5,8), in agreement with previous studies (6,7). Membrane ion channels are likely damaged following lengthening contractions and *Prox1* transfer possibly interferes with this process. It remains to be determined whether the upregulation of the membrane ion stretch-activated channel *Trpc1* induced by *Prox1* transfer in voluntary exercised *mdx* mice contributes to this improvement of excitability. However, a higher level of TRPC1 or activity of stretch-activated channels are generally associated with a worst dystrophic phenotype and fragility (39,40). In line with the present study, it was previously reported that the improved TA *mdx* muscle excitability and fragility induced by voluntary exercise and calcineurin pathway activation were also related to the changes in expression of genes encoding membrane ion channels (8).

Previous studies suggest that increased NOX2 activity is related to fragility in *mdx* mice (12,13). However, our results show that *Prox1* transfer in exercised *mdx* muscle does not reduce the expression of *Nox2* subunits (*Gp91phox, P47phox* and *Rac1*), which are shown to produce an elevated level of ROS in *mdx* mice (13). Moreover, we found no increased expression of the gene encoding the antioxidant enzyme *PrxII*, whose overexpression improves fragility in *mdx* mice (12). Finally, we found that the improvement of the fragility in response to *Prox1* transfer is not associated with significantly increased expression of *Utr* and *Des* in exercised *mdx* mice, two genes contributing to fragility in *mdx* mice (14,15).

Of note, *Prox1* transfer alone does not significantly improve fragility in sedentary *mdx* mice. The difference cannot be attributed to the fact that *Prox1* was not highly overexpressed in sedentary *mdx* mice treated with *Prox1* transfer. However, some changes induced by *Prox1* transfer are notably different between exercised and sedentary *mdx* mice: absolute maximal lengthening force (x 0.6 versus none significant change), MHC-2b (x 0.8 versus none), MHC-2x (x 1.6 versus none), *Myh7* (x 15.1 versus x 6.2), and *Trpc1* (x 2.1 versus x 1.4). Thus, our study interestingly indicates that voluntary exercise potentiates a possible gene-based therapy, at least in the preclinical field.

### *Prox1* transfer reduces maximal force production in *mdx* mice

Although *Prox1* transfer improves fragility in voluntary exercised *mdx* muscle, we found that it has a detrimental effect on absolute maximal isometric force (and maximal lengthening force). The reduced maximal strength is related to a reduced muscle weight but not either a reduction in the diameter of the different types of fibers nor a higher number of fibers of small diameter (no increase in the percentages of fibres expressing MHC-1 and MHC-2a that are smaller than the fibres expressing MHC-2b). Unexpectedly, we found that the atrophic state in response to *Prox1* transfer is associated to the downregulation of *Mstn*, a negative regulator of muscle growth in *mdx* muscle (41), without other transcriptional changes regarding several well-known atrophic processes, at least in voluntary exercised *mdx* mice. Because the relative amount of MHC-2b is reduced, we hypothesize that *Prox1* specifically reduces the synthesis of MHC-2b in exercised *mdx* mice. In line with the reduced muscle weight induced by *Prox1* transfer, several genetic or pharmacological treatments promoting slower and more oxidative fibres has been shown to induce muscle atrophy/reduced weight in *mdx* mice (26,27,29), for reasons still largely unknown.

### Conclusions

Combined to voluntary exercise, *Prox1* transfer further improves (reduced) fragility, whereas the single *Prox1* transfer approach in sedentary *mdx* mice failed to induce a significant effect on the susceptibility to exercise-induced muscle injury. Interestingly, voluntary exercise potentiates a possible gene-based therapy. This beneficial effect on fragility in exercised *mdx* mice is associated to the reduction in maximal lengthening force, the promotion of slower contractile features, and the change in *Trpc1* expression. However, *Prox1* overexpression also reduces absolute maximal force production in relation with a reduced muscle weight, despite decreased expression of *Mstn*. Thus, although Prox1 transfer aggravates at first sight muscle weakness, it can have the potential to reduce the occurrence of degeneration-regeneration cycles and consequently to slow the progression of the disease in the long term in exercised dystrophic *mdx* muscle. Is this knowledge could be exploited for therapeutic advantage?

## Acknowledgements

This study was funded by internal grants by Sorbonne Université, INSERM, Association Institut de Myologie, and Association Française contre les Myopathies awarded to AF, DF, AK, OA, and by Université de Paris awarded to AF.

We are grateful to Kari Alitalo (Wihuri Research Institute and Translational Cancer Biology Program, University of Helsinki, Finland) for the gift of Prox1 construct, Pierre Joanne (Sorbonne Université) for assistance during the experiments, Laura Julien and Sofia Benkhelifa for AAV-Prox1 production (Sorbonne Université), Delphine Bouteiller for qPCR measurements (Sorbonne Université), and Saline Jabr (Sorbonne Université) for her help in english.

